# Chromosome-level genome assembly of Lilford’s wall lizard, *Podarcis lilfordi* (Günther, 1874) from the Balearic Islands (Spain)

**DOI:** 10.1101/2023.02.15.528647

**Authors:** Jessica Gomez-Garrido, Fernando Cruz, Tyler S. Alioto, Nathalie Feiner, Tobias Uller, Marta Gut, Ignacio Sanchez Escudero, Giacomo Tavecchia, Andreu Rotger, Katherin Eliana Otalora Acevedo, Laura Baldo

## Abstract

The Mediterranean lizard *Podarcis lilfordi* is an emblematic species of the Balearic Islands. The extensive phenotypic diversity among extant isolated populations makes the species a great insular model system for eco-evolutionary studies, as well as a challenging target for conservation management plans. Here we report the first high quality chromosome-level assembly and annotation of the *P. lilfordi* genome, along with its mitogenome, based on a mixed sequencing strategy (10X Genomics linked reads, Oxford Nanopore Technologies long reads and Hi-C scaffolding) coupled with extensive transcriptomic data (Illumina and PacBio). The genome assembly (1.5 Gb) is highly contiguous (N50 = 90 Mb) and complete, with 99% of the sequence assigned to candidate chromosomal sequences and >97% gene completeness. We annotated a total of 25,663 protein-coding genes, assigning 72% to known functions. Comparison to the genome of the related species *Podarcis muralis* revealed substantial similarity in genome size, annotation metrics, repeat content, and strong collinearity, despite their evolutionary distance (~18-20 MYA). This genome expands the repertoire of available reptilian genomes and will facilitate the exploration of the molecular and evolutionary processes underlying the extraordinary phenotypic diversity of this insular species, while providing a critical resource for conservation genomics.

## 1. Introduction

*Podarcis lilfordi* (Günther, 1874), also known as Lilford’s wall lizard, is an endemic lizard of the Balearic Islands (Spain), currently confined to the Cabrera archipelago and several islets surrounding Menorca and Mallorca^1^. Due to a patchy distribution (~ 43 isolated populations) and threats from habitat loss, the species is currently listed as endangered by the IUCN (https://www.iucnredlist.org/species/17795/7481971). An extensive morphological diversity has resulted in the description of a large number (currently 23) of subspecies or morphotypes, but current understanding considers each population/islet as a distinct evolutionary unit^1^. The species exhibits several insular characteristics that likely reflect the lack of natural predators, including high population densities^2^, high survival and low fecundity^2–5^ compared to their continental relatives. The insularity and large phenotypic diversity of *P. lilfordi* have made it a popular model for studies of ecological adaptation and demographic dynamics of terrestrial vertebrates, as well as for testing predictions of how evolution proceeds on islands. During the past decades, substantial efforts have been directed toward understanding the species’ morphological diversity^1,3,6^, trophic ecology^7^, life history traits and demographic resilience^3,8^, genetics^5,9^ and associated gut microbes^10–12^. Despite this extensive knowledge and the endemic character of *P. lilfordi*, genomic information on the species has been limited^9^.

Here we present the complete genome sequence of *P. lilfordi* (both nuclear and mitochondrial) from a single female specimen (the heterogametic sex, ZW) collected from Aire Island (Menorca), where the species was first described (Gunther, 1874). We used a sequencing strategy combining long and short reads, coupled with RNAseq data from multiple tissues and specimens, to achieve a highly robust assembly and annotation. The *P. lilfordi* genome, along with that of its continental relative *Podarcis muralis*^13^, will extend current genomic resources within this highly diverse lizard genus^14,15^ and provide a genetic framework to begin understanding the processes behind the remarkable diversification of the Lilford’s wall lizard. In addition, the genome will provide a critical tool for the development of a conservation plan based on population-level genomics (accounting for levels of genetic diversity, drift, and accumulation of deleterious mutations) for an accurate assessment of the species adaptive potential and resilience in face of current challenges by global change and increasing human pressure in the Balearic Islands.

## 2. Materials and Methods

### 2.1. Sample collection and processing

Five adult specimens of *P. lilfordi* (*subspecies lilfordi*) were collected in April 2021 on Aire Island, to the southeast of Menorca Island (Spain). The island is one of the largest and most densely populated (surface area 34 ha, 4098.60 lizards/ha)^4^. Specimens were caught using pitfall traps containing fruit juices placed along paths and vegetation edges. All specimens were sexed according to visual examination of femoral pores and morphology^16^, weighed, and body size measured as snout to vent length (SVL) (see Supplementary Table S1 for metadata). Individuals were sacrificed by cervical dislocation, immediately stored on dry ice, and kept at −80°C until processing. Under a sterile hood, each frozen specimen was rapidly dissected to extract all major organs, including heart, liver, lungs, kidneys, brain, testicles/ovaries, intestine, and muscle tissue from the tail. All tissues were immediately stored at −80°C.

A single female adult specimen (the heterogametic sex, ZW) was chosen for genome sequencing. Tissues were sent to the Centre Nacional d’Anàlisi Genòmica (CNAG-CRG) for DNA extraction and sequencing. Samples from the remaining specimens were shipped to Lund University (Sweden) for RNA extraction. RNA sequencing was performed at the SNP&SEQ Technology Platform (for short reads), and at the Uppsala Genome Center (for long reads) (SciLifeLabs, Sweden).

Specimen sampling was granted by the Species Protection Service (Department of Agriculture, Environment and Territory, Government of the Balearic Islands), under permit CAP03/2021 (to LB). The specimen remains were deposited at the Museum of Natural History of Barcelona (Spain), under voucher name MZB 2022-5701.

### 2.2. Genomic DNA extractions

High-molecular-weight (HMW) gDNA was extracted from frozen liver using the Nanobind tissue kit (Circulomics) following the manufacturer’s protocol. Briefly, two cryopreserved liver aliquots of 38 mg and 24 mg were homogenized under cryogenic conditions on dry ice, using mortar and pestle. The pulverized tissue was collected into 1.5 ml tubes with lysis buffer (Circulomics). Nanobind disk (Circulomics) was used on fresh supernatant for the gDNA binding. The HMW gDNA eluate was quantified by Qubit DNA BR Assay kit (Thermo Fisher Scientific) and the DNA purity was evaluated using Nanodrop 2000 (Thermo Fisher Scientific) UV/Vis measurements. The gDNA integrity was evaluated with pulsed-field gel electrophoresis SeaKem^®^ GOLD Agarose 1% (Lonza), using the Pippin Pulse (Sage Science). The gDNA samples were stored at 4 °C.

### 2.3. Genome sequencing

The complete genome sequence was achieved employing a mixed sequencing strategy, combining the use of 10X linked short reads (Illumina NovaSeq 6000, 2×150bp) for base accuracy, long reads (Oxford Nanopore Technologies, ONT) for high contiguity and repeat resolution, and Hi-C (Illumina NovaSeq 6000, 2×150bp) for chromosome-level scaffolding (Fig. 1).

**Figure 1:**
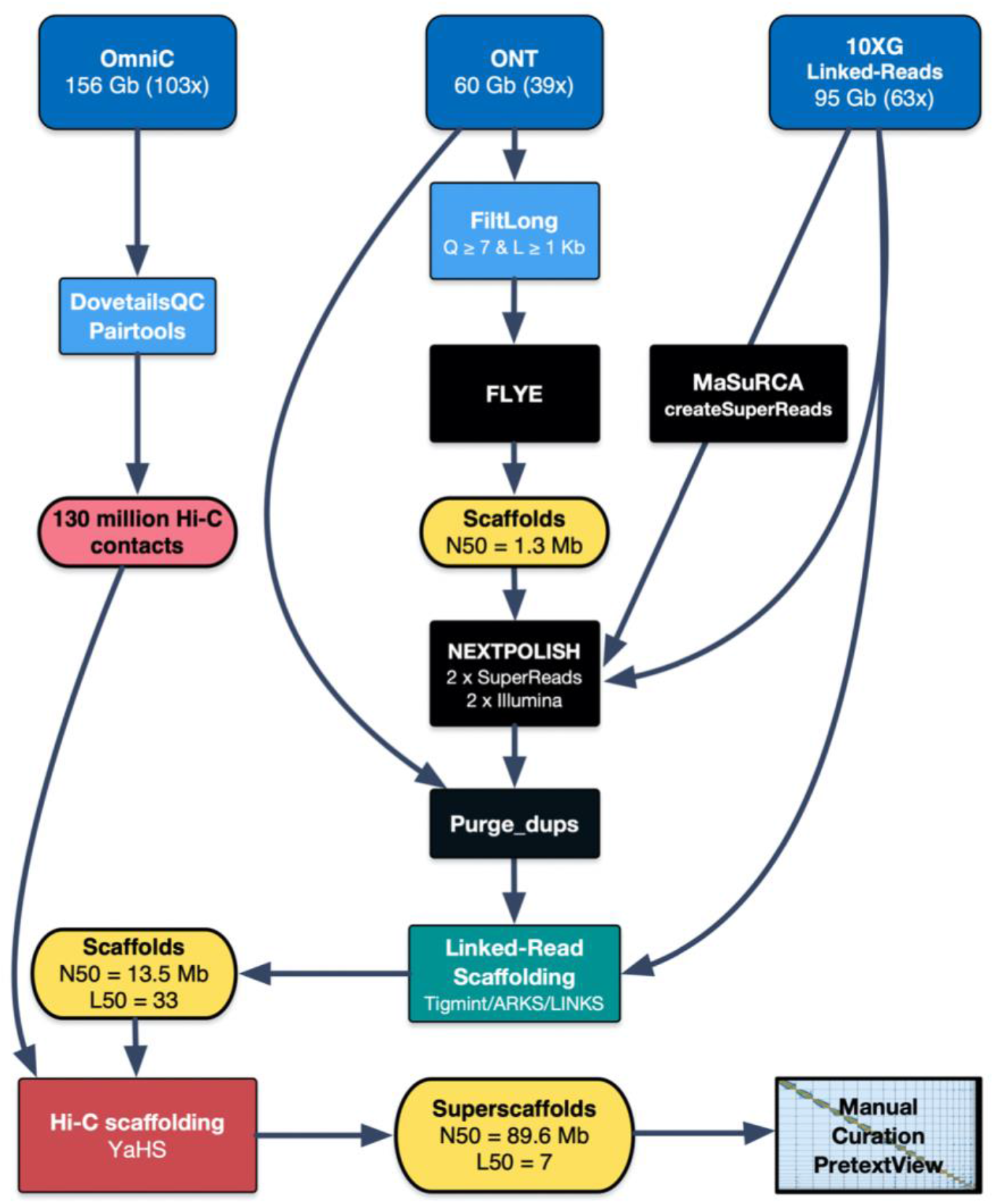
Assembly Workflow. Summary of the main steps followed to obtain the curated chromosome-level assembly. See the Snakemake pipeline for detailed information (https://github.com/cnag-aat/assembly_pipeline).

The linked reads library was prepared using the Chromium Controller instrument (10X Genomics) and Genome Reagent Kits v2 (10X Genomics) following the manufacturer’s protocol. Briefly, 10 ng of HMW gDNA was portioned in GEM reactions including unique barcode (Gemcode) after loading onto a chromium controller chip. The droplets were then recovered, isothermally incubated, fractured and the intermediate DNA library then purified and size-selected using Silane and Solid Phase reverse immobilisation (SPRI) beads. Illumina-compatible paired-end sequencing libraries were prepared following 10X Genomics recommendations and validated on an Agilent 2100 BioAnalyzer with the DNA 7500 assay (Agilent). The library was sequenced on NovaSeq 6000 (Illumina, 2×151bp) following the manufacturer’s protocol for dual indexing. Image analysis, base calling and quality scoring of the run were processed using the manufacturer’s software Real Time Analysis (RTA v3.4.4) and followed by generation of FASTQ files.

The ONT libraries were prepared using the 1D Sequencing kit SQK-LSK110. Briefly, 1.0 μg of the HMW gDNA was DNA-repaired and DNA-end-repaired using NEBNext FFPE DNA Repair Mix and the NEBNext UltraII End Repair/dA-Tailing Module (New England Biolabs, NEB), followed by ligation of sequencing adaptors. The library was purified with 0.4 X AMPure XP Beads and eluted in Elution Buffer. Four sequencing runs were performed on GridION Mk1 (ONT) using an R9.4.1 flow cell (FLO-MIN106D, ONT) on the MinKNOW platform version 4.2.5 for real-time monitoring. Data was collected for 110 hours and basecalled with Guppy version 4.3.4 in high accuracy mode using the dna_r9.4.1_450bps_modbases_dam-dcm-cpg_hac.cfg model.

The Hi-C libraries were prepared using the Omni-C kit (Dovetail Genomics), following the manufacturer’s protocol. Briefly, frozen muscle tissue (heart) was pulverised using mortar and pestle immersed in a liquid nitrogen bath. Chromatin was fixed in place with formaldehyde (Sigma Aldrich), digested with DNase I and DNA extracted. DNA ends were repaired, and a biotinylated bridge adapter was ligated followed by proximity ligation of adapter-containing ends. After reverse crosslinking, the DNA was purified and followed by preparation of Illumina-compatible paired-end sequencing libraries (omitting the fragmentation step). Biotinylated chimeric molecules were isolated using streptavidin beads before PCR enrichment of the library. The library was sequenced on NovaSeq 6000 (Illumina, 2×151bp) following the manufacturer’s protocol for dual indexing.

### 2.4. Transcriptome sequencing

RNA was extracted from five different tissues (heart, kidney, liver, lungs, and tail) of five specimens using the RNeasy Mini Kits (Qiagen) and including an on-column DNA digestion step. RNA concentration was measured using the Qubit RNA High Sensitivity assay and RNA quality was assessed using the Eukaryote Total RNA Nano assay on the Agilent 2100 Bioanalyzer system. For short-read RNA sequencing, we pooled samples of 2-4 individuals per tissue in equimolar amounts (Supplementary Table S2). These five tissue-specific samples were subjected to Illumina’s TruSeq Stranded mRNA library preparation kit and sequenced on NovaSeq 6000 (Illumina, 2×150 bp). For long-read RNA sequencing, we pooled five individual tissue samples in equimolar amounts (Supplementary Table S2) and subjected this single, mixed sample to PacBio Iso-Seq sequencing.

### 2.5. Nuclear genome assembly

The nuclear genome was assembled following the workflow summarised in Fig. 1. To increase reproducibility, the main steps have been included into a snakemake^17^ pipeline (https://github.com/cnag-aat/assembly_pipeline). A few modifications were added for this project which are detailed below, along with a summary of major steps. All intermediate and final assemblies produced during the process were evaluated as indicated in the pipeline description, using BUSCO^18,19^ v 5.4.0 with vertebrata_odb10, fasta-stats.py (https://github.com/galaxyproject/tools-iuc), Nseries.pl and Merqury^20^ v1.1.

Before assembly, the 10X Illumina reads were preprocessed using LongRanger Basic v2.2.2 and the nanopore reads filtered using FiltLong v0.2.0 with options ‘--min_length 1000 --min_mean_q 80’. The resulting set of filtered nanopore reads accounted for 50.8 Gb (about 34x coverage) with a mean Phred-scale read quality of 9.8 and a read length N50 of 31.4 Kb (Supplementary Table S3).

First, we assembled the filtered nanopore reads using Flye v2.8.3 with options ‘--nano-raw -i 2’ (the latest performs two final rounds of polishing in the output assembly) obtaining a base assembly that comprised a total of 1.54 Gb, with an N50 of 1.36 Mb and a consensus quality (QV) of 27.19 (Fig. 1 and Supplementary Table S4).

Second, to maximise the QV of the Flye assembly, we tested several polishing tools and strategies (Supplementary Table S4). The best results were obtained using a combined strategy that exclusively used 10X illumina reads to improve the QV. We used MaSuRCA^22,23^ version 4.0.4 using *createSuperReadsForDirectory.perl* to obtain high-quality SuperReads with a read N50 of 539 bp. The SuperReads were then aligned to the genome with Minimap2^24^ v2.17 and the Illumina reads were aligned with BWA-MEM^25^ v0.7.15. Finally, we ran NextPolish^26^ v1.3.1 with these SuperReads, treating them as long reads, and the debarcoded 10X linked reads as paired-end Illumina for polishing the Flye assembly. In total, four polishing iterations were performed with NextPolish: two with the SuperReads and another two with the Illumina reads.

Third, to obtain a haploid reference (removing alternate haplotigs and other artificial duplications), we purged the assembly using purge_dups^27^ v1.2.5 with cutoffs -l 5 -m 24 -u 78. This step removed 4,289 scaffolds accounting for 94,919,135 bp.

Fourth, the purged assembly was corrected and scaffolded with 10X Linked-Reads using the Faircloth’s Lab pipeline (http://protocols.faircloth-lab.org/en/latest/protocols-computer/assembly/assembly-scaffolding-with-arks-and-links.html#). This pipeline includes Tigmint^28^ v1.1.2, ARKS^29^ v1.0.3, and LINKS^30^ v1.8.5. Tigmint was used to identify and correct mis-assemblies using the linked reads in the purged assembly. The corrected assembly was then scaffolded using ARKS and LINKS, resulting in 2,783 scaffolds accounting for 1,460,390,273 bp with an N50=13.51 Mb (Supplementary Table S4).

Finally, the Omni-C reads were then mapped to the assembly using BWA-MEM and pre-processed using the Dovetail pipeline (https://omni-c.readthedocs.io/en/latest/fastq_to_bam.html). The parsing of the read pairs was done with the default minimum mapping quality of 40. After removal of PCR duplicates (46.34%, see Supplementary Table S5), 130,208,672 read pairs remained and were used as input to YaHS^31^ v1.1 scaffolder with default parameters. We performed two rounds of assembly error correction and made 15 breaks, followed by ten rounds of scaffolding from higher to lower resolution (10 Mb down to 10 Kb), which produced an assembly with a span of 1,460,440,873 bp with an N50 of 89.49 Mb. The high contiguity achieved is also reflected in the scaffold L90 of 17, a value close to the number of chromosomes (n=19) (Fig. 1 and Supplementary Table S4).

### 2.6. Manual Curation

To guide manual curation of the assembly, we computed the mean Illumina coverage for all scaffolds (using BWA-MEM v0.7.15, SAMtools^32^ v1.9 and BlobTools^33^ v1.1) and the whole-genome alignments (WGA) against the genome assemblies of two species belonging to family *Lacertidae:* a ZZ male of *Podarcis muralis* (PodMur1.0) and a ZW female of *Lacerta agilis* (rLacAgi1.pri). The WGA alignments were produced with nucmer4^34^ and visualised with Dot (https://github.com/MariaNattestad/dot). This approach allowed us to “scaffotype” (i.e., assign to a chromosome) the largest 20 superscaffolds (Supplementary Table S6 and S7). Finally, the location of gaps (fasta-stats.py) and telomeres (https://github.com/tolkit/telomeric-identifier), together with the Illumina coverage, were added to the contact map using PretextGraph (https://github.com/wtsi-hpag/PretextGraph). Manual curation was then performed using PretextView (https://github.com/wtsi-hpag/PretextView). Given the high quality and contiguity of the YaHS assembly, the curation involved only 17 edits.

The Blobtoolkit^35^ pipeline was then run on the curated assembly (Supplementary Fig. S1), using the NCBI nt database (updated on September 2022) and several BUSCO odb10 databases (sauropsida, vertebrata, metazoa, eukaryota, chlorophyta, fungi and bacteria). Removal of six contaminated scaffolds (based on GC cutoff of 0.3 - 0.65) (see Supplementary Table S8) resulted in the final assembly rPodLil1.2.

Finally, by comparing Illumina and ONT coverage estimates, we calculated the ratio of coverage for each sex chromosome with respect to the autosomal mean coverage (Supplementary Table S9). Whole genome alignments of the *P. lilfordi* (rPodLil1.2) against *P. muralis* (PodMur1.0) were performed with Minimap2 using the ‘-x asm5’ option and visualized with the *pafr* R package.

### 2.7. Nuclear genome annotation

Gene annotation was achieved by combining transcript alignments, protein alignments and *ab initio* gene predictions (see flowchart in Supplementary Fig. S2).

Prior to gene annotation, repeats were searched with RepeatMasker v4-1-2 (http://www.repeatmasker.org) using the custom RepBase repeat library available for *Podarcis*, along with a specific repeat library generated with RepeatModeler^36^ v1.0.11 for our assembly. After excluding those repeats that were part of repetitive protein families (performing a BLAST search against Uniprot; last accessed March 2022), a final repeat annotation was produced using RepeatMasker. As this repeat annotation was produced mainly to aid in the genome annotation, low-complexity repeats were not annotated. The *P. muralis* genome was similarly processed for repeat annotation and comparative analysis.

Long and short RNA reads were aligned to the genome assembly using STAR^37^ v-2.7.2a and Minimap2 v2.14 (with ‘-x splice:hq-uf’ options), respectively. Transcript models were subsequently generated using StringTie^38^ v2.1.4 on each BAM file and then all the models combined using TACO^39^ v0.6.3 were given as input for the Program to Assemble Spliced Alignments (PASA^40^ v2.4.1) to produce PASA assemblies for annotation. High-quality junctions used during the annotation process were obtained with Portcullis^41^ v1.2.0 after STAR and Minimap2 mapping. Finally, to detect coding regions in the transcripts, the *TransDecoder* program (embedded in the PASA package) was run on the PASA assemblies. For protein assignment, the complete proteomes of *P. muralis, Pogona vitticeps* and *Pantherophis guttatus* were downloaded from Uniprot in April 2022 and aligned to the genome using Spaln^42^ v2.4.03. *Ab initio* gene predictions were performed on the repeat-masked rPodLil1.1 assembly with three different programs: GeneID^43^ v1.4, Augustus^44^ v3.3.4 and Genemark-ES^45^ v2.3e, with and without incorporating evidence from the RNAseqdata. The gene predictors were run with trained parameters on humans except for Genemark, which runs in a self-trained mode. Finally, all the data were combined into consensus CDS models using EvidenceModeler-1.1.1^40^ (EVM). Additionally, untranslated regions (UTRs) and alternative splicing forms were annotated via two rounds of PASA annotation updates (Supplementary Fig. S2).

Functional annotation was performed on the annotated proteins with Blast2GO^46^. First, a DIAMOND Blastp^47^ search was made against the NCBI nr database (last accessed May 2022). Furthermore, InterProScan^48^ was run to detect protein domains on the annotated proteins. All these data were combined by Blast2GO, which produced the final functional annotation.

General statistics on the genome and individual chromosomes were computed with in-house PERL scripts. The sex chromosome W presented 14 transposon-derived, short, only *ab-initio* single-copy genes. Due to the high repetitiveness of the chromosome, these genes were considered as artefacts and removed from the final annotation.

The annotation of non-coding RNAs (ncRNAs) was obtained as follows. First, the program CMsearch^49^ v1.1 from the Infernal^50^ package was run against the RFAM database of RNA families v12.0. Additionally, transfer RNA genes were identified by tRNAscan-SE^51^ v2.08. Long non-coding RNAs (lncRNAs) were identified as those expressed transcripts (assembled by PASA) longer than 200 bp that were not included in the protein-coding annotation and not covered by a small ncRNA in more than 80% of their length. The resulting transcripts were clustered into genes (i.e., same gene assignment) using shared splice sites or significant sequence overlap.

### 2.8. Mitogenome assembly and annotation

To obtain the mitochondrial sequences, all ONT reads, previously filtered for whole-genome assembly with FiltLong v.0.2.0 to be at least 1 Kb long and have a mean quality of 7, were mapped with Minimap2 against the *P. muralis* complete mitochondrial genome (NC_011607.1; 17,311 bp) with options: ‘-t $THREADS - ax map-ont $DATABASE $READS’. We retained all reads with mapping quality = 12 (relatively unique) and at least 800 exact matches to the mitochondrial genome reference; these included 8,093 reads and a total of 51,692,448 bp (estimated mitochondrial coverage 2,986x).

All filtered ONT reads were assembled with Flye^21^ v2.9 using the options: ‘flye --meta --scaffold -t 12 -i 2 -g 25k --nano-raw’. The ‘—meta’ option is the most appropriate for uneven coverage samples and two polishing iterations were run with the ONT reads on the final assembly with ‘-i 2’. Finally, the output assembly was screened for circular contigs, resulting in eight linear contigs and a single circular contig 17,112 bp long (contig_1).

As the reference mitogenome (*P. muralis*) is relatively distant (18-20 MYA), the identification of Illumina reads mapping outside the most conserved regions of the organelle is not straightforward. To overcome this issue, we mapped all the Illumina data (previously de-barcoded PE 2×150bp reads 10x linked-reads) to our complete Flye long-read assembly with gem-mapper, with ≤2% mismatches. Finally, a total of 326,105 read pairs were collected for further polishing (estimated coverage of the mitochondrial genome is 5,255.81x).

To further improve the sequence accuracy of the assembled mitochondrial genome, we performed two additional rounds of polishing on contig_1 with the selected Illumina reads using NextPolish v1.1.0 with Illumina PE 2×150bp with options: ‘-paired -max_depth 5000’. The polished assembly was evaluated with Merqury v1.1 using ‘k=21’ on the mitochondrial Illumina reads, *dnadiff* from MUMmer^34^ package 4.0.0beta2 and fasta-stats.py. The resulting circular chromosome was rotated and oriented according to the *P. muralis* reference, after detecting the appropriate Origin coordinates with the *dnadiff*.

The annotation of the mitogenome was performed using the MITOS^53^ Web Server (http://mitos.bioinf.uni-leipzig.de/). Manual curation was performed by checking the predicted sequences and comparing the annotation to that of the *P. muralis* mitogenome. As a result, one partial tRNA-Asp was removed from the annotation due to its absence in the *P. murallis* mitogenome annotation and to the high e-value reported by MITOS (e-value = 0.04071).

## 3. Results and Discussion

### 3.1. Genome assembly

Genome sequencing yielded a total of 95 Gb of Illumina data (2×150 bp), 156.2 Gb of Omni-C data (2×150 bp) and 60 Gb of ONT data (Fig. 1). Assembly of the ONT data, followed by polishing, scaffolding, and manual curation resulted in a highly contiguous and complete assembly (rPodLil1.2) of 1.46 Gb, in line with the *C-value* and assembly span of *P. muralis* (1.51 Gb for PodMur1.0)^13^ (Fig. 2). It has a contig N50 of 1.48 Mb, scaffold N50 of 89.64 Mb (≥10Mb) and QV of 40 (Table S4), meeting the minimum quality requirement of 6.C.Q40 (megabase contig N50 and chromosomal-scale scaffold N50, with less than 1/10,000 error rate) established by the Earth Biogenome Project (EBP) for eukaryotic species with sufficient DNA and tissue^55^.

**Figure 2:**
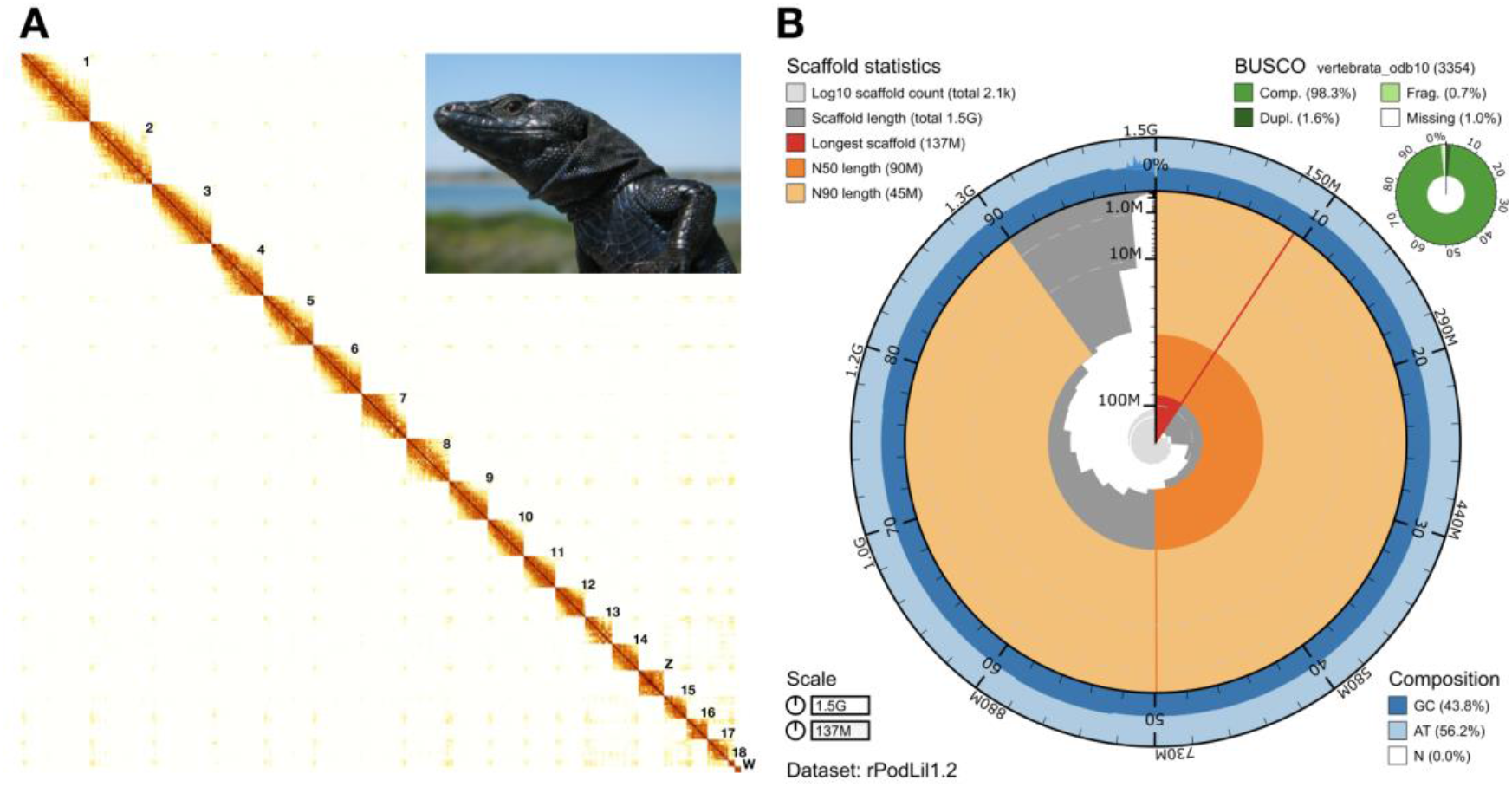
Visual summary of the rPodLil1.2 assembly. (A) Hi-C contact map of the genome assembly visualised in PretextView. The map shows 20 superscaffolds, ordered from longest to shortest, corresponding to the 18 autosomes and the two sex chromosomes (Z and W); the latter can be better visualized in Fig. S4. Additionally, there are a total of 44 short unlocalized scaffolds within the main chromosomes, and 2,084 short unplaced scaffolds (not visible at the resolution shown). (B) Snailplot summarising assembly metrics, including scaffold statistics, BUSCO completeness, total size, and base composition. The main plot is divided into 1,000 size-ordered bins around the circumference with each bin representing 0.1% of the 1.460.085.851 bp assembly. The distribution of record lengths is shown in dark grey with the plot radius scaled to the longest record present in the assembly (137 Mb, shown in red). Orange and pale-orange arcs show the N50 and N90 record lengths (90 Mb and 45 Mb), respectively. The pale grey spiral shows the cumulative record count on a log scale with white scale lines showing successive orders of magnitude. The blue and pale-blue area around the outside of the plot shows the distribution of GC, AT, and N percentages in the same bins as the inner plot. A summary of complete, fragmented, duplicated, and missing BUSCO genes in the vertebrata_odb10 set is shown in the top right.

It is consistent with the karyotype (2n=38)^54^ with 98.70% of the sequence assigned to candidate chromosomal sequences, 18 autosomes and two sex chromosomes (Fig. 2). Moreover, it is complete, with 98.3% of single copy complete genes and a k-mer completeness of 87%, and a low false duplication rate of 0.68%.

Assignment to chromosomes was done via whole-genome alignments to the *P. muralis* (a male) and *L. agilis* (rLacAgi1.pri, a female) genome assemblies. The alignments revealed a high level of collinearity to both assemblies for the autosomes as well as the Z chromosome (shown for *P. muralis* in Fig S4A). Sex chromosome assignment was additionally supported by manual curation, showing the expected drop in sequencing coverage (Table S9) and typical pattern of Hi-C contacts (Fig. S4). Sex chromosomes in lacertids (ZZ and ZW) are known to differ in gene copy numbers, with males (ZZ) showing twice as many genes as females (ZW) due to W degeneration^55^. Moreover, the W is mostly heterochromatic and shows a typical high content in repetitive sequences^56,57^, reducing the number of informative Hi-C read pairs for scaffolding in this chromosome. As expected, the Z chromosome of *P. lilfordi* was assembled into one superscaffold of 50.7 Mb in size, while the W chromosome was partially assembled into a 12.3 Mb superscaffold that aligns to the superscaffold W of *L. agilis* (Fig. S4C).

### 3.2. Genome annotation

We annotated a total of 25,663 protein-coding genes that produce 43,578 transcripts (1.7 transcripts per gene) encoding 38,615 unique protein products (Table 1). We were able to assign functional labels to 72% (29,273) of the annotated proteins. The annotated transcripts contain 11 exons on average, with 91% of them being multi-exonic. In addition, we annotated 47,052 non-coding transcripts, including 12,785 lncRNAs and 34,267 sncRNAs (Table 1).

**Table 1:** Final genome annotation statistics for the *P. lilfordi* assembly (rPodLil1.2) and comparison to *P. muralis* (PodMur1.0)

|  | **rPodLil1.2** | **PodMur1.0** |
| --- | --- | --- |
| Genome size (bp) | 1,460,085,851 | 1,511,002,858 |
| Number of protein-coding genes | 25,663 | 22,062 |
| Median gene length (bp) | 13,456 | 11,808 |
| Number of transcripts | 43,578 | 37,240 |
| Number of proteins | 38,615 | 36,445 |
| Proteins with complete ORFs | 37,249 | 30,503 |
| Functionally annotated proteins | 29,273 | 25,768 |
| Number of coding exons | 232,494 | 219,498 |
| Median UTR length (bp) | 2,138 | 505 |
| Median intron length (bp) | 1,297 | 1,245 |
| Exons/transcript | 10.90 | 12.12 |
| Transcripts/gene | 1.70 | 1.69 |
| Multi-exonic transcripts | 0.91 | 0.92 |
| Gene density (gene/Mb) | 17.58 | 14.60 |

Running BUSCO on both the annotated protein-coding transcripts and protein sets showed a gene completeness of 97.9% and 97.1%, respectively, using the vertebrata_odb10 database. These results are in line with the genome BUSCO completeness of 98.3% and demonstrate the high accuracy of the genome annotation pipeline. Minor differences in the results obtained using the three sources of evidence are likely due to the algorithm that BUSCO uses, and the threshold established to consider a gene absent, fragmented or complete.

Comparison of *P. lilfordi* and *P. muralis* genome annotation statistics show only few differences (Table 1). Gene content is comparable, although we annotated ~3,000 extra protein-coding genes for *P. lilfordi*. Alignment of these genes back to the PodMur1.0 genome assembly indicates that only 600 are unique to *P. lilfordi* (i.e., do not align to the *P. muralis* assembly). Most of the *P. lilfordi* proteins (96.5%) have a complete Open Reading Frame (ORF) (versus 83.7% in *P. muralis*) supporting our high-quality assembly and annotation (Table 1). Moreover, median UTR length is four times longer in *P. lilfordi* than in *P. muralis*, which is likely due to the usage of long-read transcript sequencing technologies (PacBio) for our annotation. Although most of the observed differences can be ascribed to the use of different evidence sources and annotation pipelines, additional genome resequencing data will be critical for validation.

### 3.3. Repetitive elements

Around 39% of the assembled genome was repetitive: 6% was annotated as repeats using the “podarcis” Repbase library and an additional 33% was classified as repeats using the repeat library obtained after running Repeat Modeler (https://www.repeatmasker.org/RepeatModeler/) against the assembly (Table 2). These repeats were represented by transposable elements, including short, interspersed elements (SINEs), long interspersed nuclear elements (LINEs), long terminal repeat retrotransposons (LTRs), and DNA transposons. The repeat content landscape of *P. lilfordi* was highly comparable to that of *P. muralis* (38-39%), in line with their similar genome size^58^: most of the repeats were classified as LINEs (~12%), followed by DNA transposons (~7%) and SINEs (4-5%) (Table 2). Percentage of LTR was slightly higher in *lilfordi* than *muralis* (2.37% against 1.36%). Overall, we did not detect any major differences in repeat content between the two *Podarcis* species.

**Table 2:**
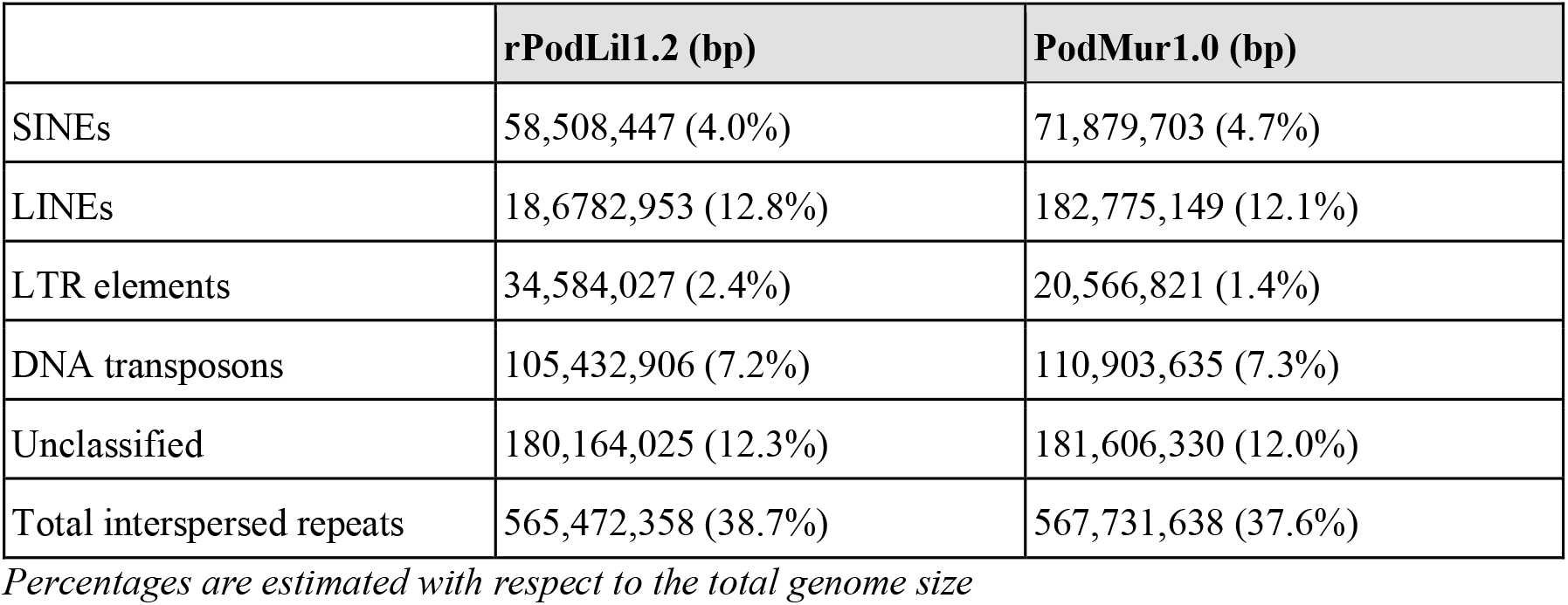
Repeat content of the *P. lilfordi* assembly and comparison with *P. muralis*.

### 3.4. Mitogenome

The resulting mitogenome assembly is a single circular contig 17,251 bp long with a QV of 44.12. Its GC content is 38.73%, very similar to the 38.55% observed in the *P. muralis* mitogenome. As expected, the average identity between both mitogenomes is 88.74%. The origin of replication was identified and used to linearize the circular chromosome by comparison to the *P. muralis* mitogenome. After annotation it was confirmed that this was the origin of the tRNA-Phe (MT -TF) that is used as a standard origin in vertebrate mitogenomes^59^. The final annotation of the mitochondrial chromosome contained 13 protein-coding genes, 2 rRNAs and 22 tRNAs, in line with expectations for vertebrates.

## 4. Conclusions

Here we announce and publicly release the *Podarcis lilfordi* reference nuclear and mitochondrial genome assembly. This is the first chromosome-level reference for an endemic reptile species released within the framework of the Catalan Initiative for the Earth Biogenome Project (CBP). The *P. lilfordi* genome is the second complete genome within the highly diverse genus *Podarcis*, along with that of *P. muralis*. Despite their evolutionary divergence (18-20 MYA^14^), the two species did not reveal any major structural difference, suggesting a substantial conservation in genome organisation and overall annotation. Given its high quality, contiguity and annotation, the *P. lilfordi* genome sequence will represent a valuable resource for evolutionary and conservation genomics studies. The resource will facilitate comparative genomics of Lacertidae and reptiles in general, and aid in the understanding of the genetic bases of vertebrate insular adaptation (i.e., the island syndrome) and demographic resilience. Additionally, the genome will represent a critical reference to explore the genetic diversity of this endemic species, its adaptive plasticity and local adaptation, along with its ability to respond to current threats by human pressure in the Balearic Islands. Finally, we expect that future genome analyses will have a critical impact on conservation management and policy decisions on this endangered species.

## Acknowledgments

We would like to thank the following people and institutions: The Catalan Initiative for the Earth Biogenome Project (CBP) for coordination and promotion efforts of this genome project; Yumi Sims, Jo Wood and Alan Tracey (Wellcome Sanger Institute) for their help and advice on assembly curation; Marc Palmada-Flores (IBE-UPF) for his feedback on assembly and advice about available lizard genomes; Chenxi Zhou (Wellcome Sanger Institute) for his help on producing contact maps in different formats directly from the YaHS output; Emiliano Trucchi (Marche Polytechnic University, Italy) and Giorgio Bertorelle (University of Ferrara, Italy) for access to the draft *Podarcis raffonei* chromosome W sequence (unpublished at the time of this study). Finally, we are also grateful to Dovetail Genomics (Cantata Bio) for their support with the quality control and pre-processing of Omni-C data before scaffolding.

## Author contributions

L.B., G.T., N.F. and T.U. conceived and advised the project. L.B and K.E.O.A. performed sample collection and preparation. M.G, I.S.E. and N.F performed data production. J.G.G, F.C. and T.S.A performed data processing and genome assembly. J.G.G and T.S.A performed data processing and genome annotation. All authors wrote and approved the final manuscript.

## Data availability

Genome sequencing data have been submitted to the European Nucleotide Archive (ENA) (https://www.ebi.ac.uk/ena/browser/home) under accession number PRJEB50058. The genome assembly and annotation have also been deposited to the ENA under the accession number GCA_947686815.1, as part of project PRJEB47961. The transcriptomic data produced for genome annotation have been deposited in the National Center for Biotechnology Information (NCBI) under the Bioproject accession ID PRJNA897120. For convenience, all the above data are collected under the ENA umbrella study PRJEB50294. In addition, a genome browser, a BLAST Sequence Server and annotation data can be found at https://denovo.cnag.cat/podarcis. A Snakemake pipeline for genome assembly is available at https://github.com/cnag-aat/assembly_pipeline.

## Supplementary data

**Table S1:** Sampled specimen information.

**Table S2:** RNA samples and quality.

**Table S3:** ONT read statistics.

**Table S4:** Assembly statistics, including intermediate steps.

**Table S5:** Omni-C mapping statistics.

**Table S6:** Scaffotyping based on Whole Genome Alignments (Yash vs PodMur1.0).

**Table S7:** Scaffotyping based on Whole Genome Alignments (Yash vs rLacAgi1).

**Table S8:** Contaminated scaffolds.

**Table S9:** Ratio of sex chromosome coverage with respect to the autosomes mean coverage (based on ONT and Illumina reads).

**Figure S1: Hexagon-binned blob plot of base coverage of filtered ONT reads against GC proportion for scaffolds in assembly rPodLil1.1.** Scaffolds are coloured by phylum (according to Blast hits against the nt database; last accessed September 2022) and binned at a resolution of 30 divisions on each axis. Coloured hexagons within each bin are sized in proportion to the sum of individual scaffold lengths on a square-root scale, ranging from 1,018 to 693,200,870. Histograms show the distribution of scaffold length sum along each axis.

**Figure S2: Annotation flowchart based on the non-decontaminated assembly (rPodLil1.1).**

**Figure S3: K-mer comparison between the Illumina reads and the rPodLil1.2 assembly.** Stacked histogram of k-mer distributions obtained by comparing the assembly with Merqury v1.1 using k=21 on the 10X Illumina reads. Artificial duplications corresponding to duplicate k-mers are shown in blue above the main peak (~40x). They only account for 0.68% of the k-mers.

**Figure S4: Chromosome assignment.** (A) Whole genome alignment of *P. lilfordi* to *P. muralis*. Chromosomal sequences, named according to corresponding chromosomal sequences in *P. muralis*, are ordered from largest to smallest in *P. lilfordi* and oriented with respect to *P. muralis*, which inverts the order of chromosomes 16 and 17. Alignments longer than 100 kb were selected and visualised with the *pafr* R library. (B) Hi-C contact map showing scaffolds corresponding to the sexual chromosomes. (C) Alignment of the scaffold assigned to the W chromosome in *L. agilis* against the corresponding scaffold in *P. lilfordi*.

