## Supplementary Fig. for "Chromosome-level genome assembly of Lilford’s wall lizard, *Podarcis lilfordi* (Günther, 1874) from the Balearic Islands (Spain)"

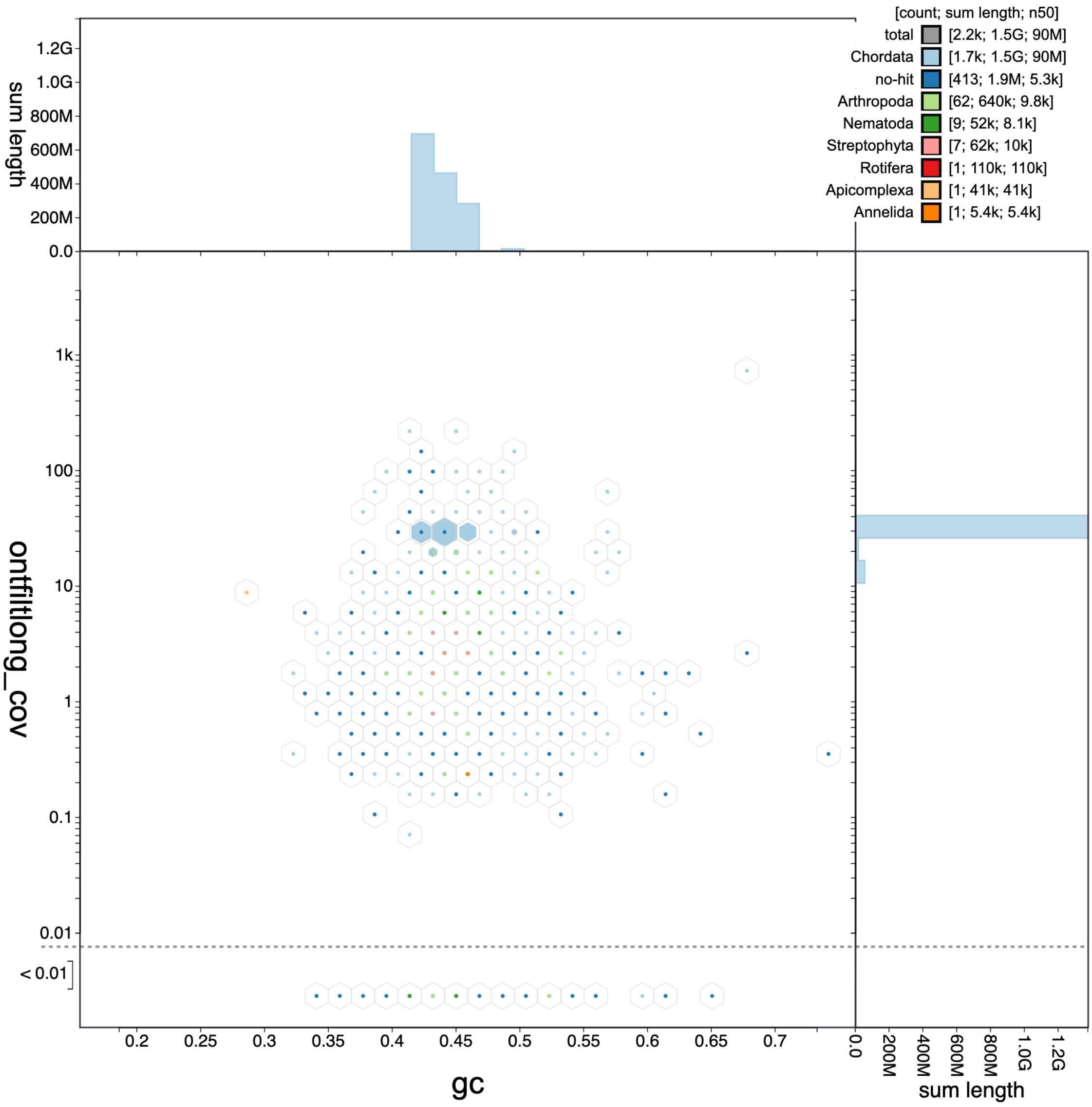


**Figure S1: Hexagon-binned blob plot of base coverage of filtered ONT reads against GC proportion for scaffolds in assembly rPodLil1_1.** Scaffolds are coloured by phylum (according to Blast hits against the nt database; last accessed September 2022) and binned at a resolution of 30 divisions on each axis. Coloured hexagons within each bin are sized in proportion to the sum of individual scaffold lengths on a square-root scale, ranging from 1,018 to 693,200,870. Histograms show the distribution of scaffold length sum along each axis.

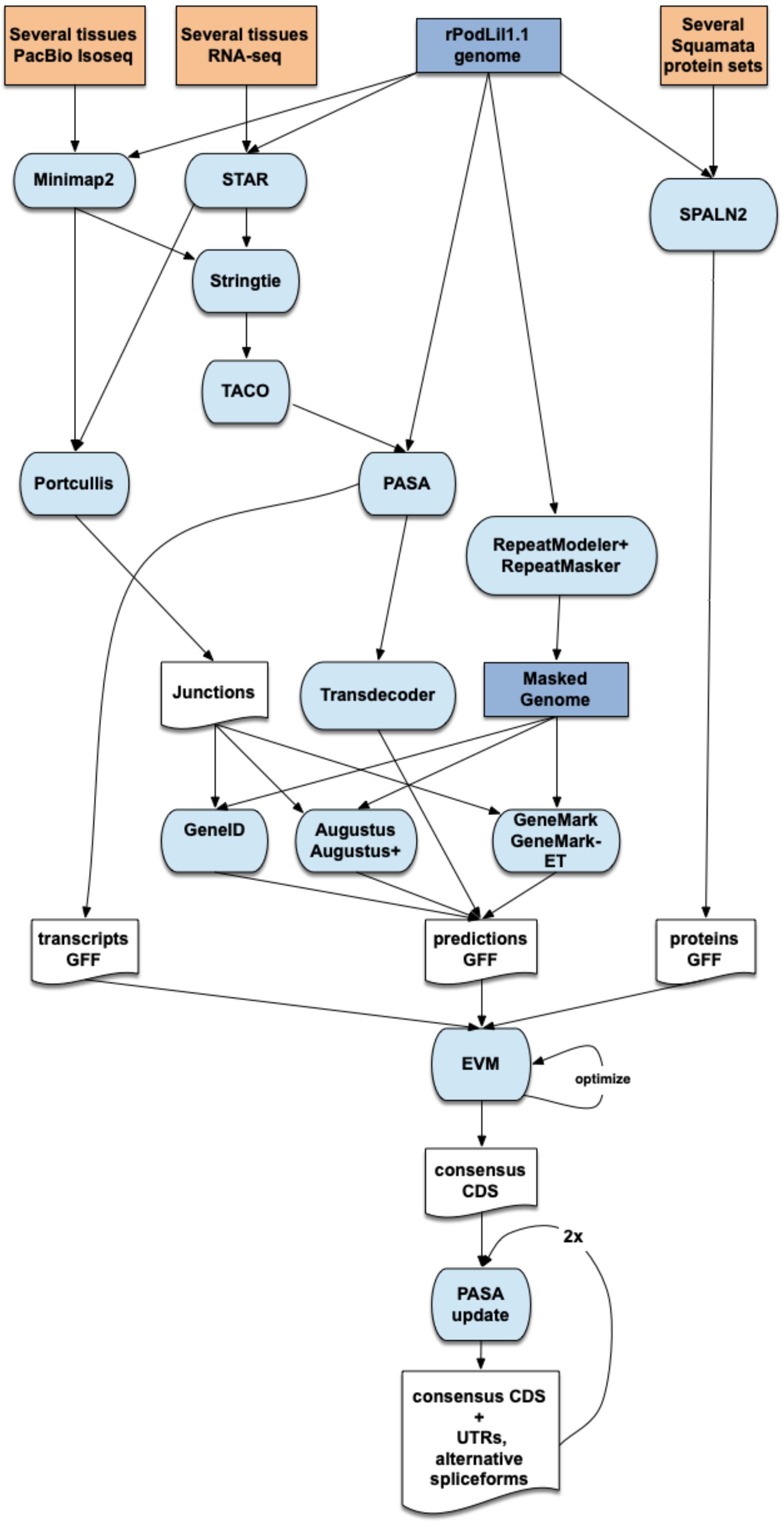


**Figure S2: Annotation flowchart based on the non-decontaminated assembly (rPodLil1.1).**

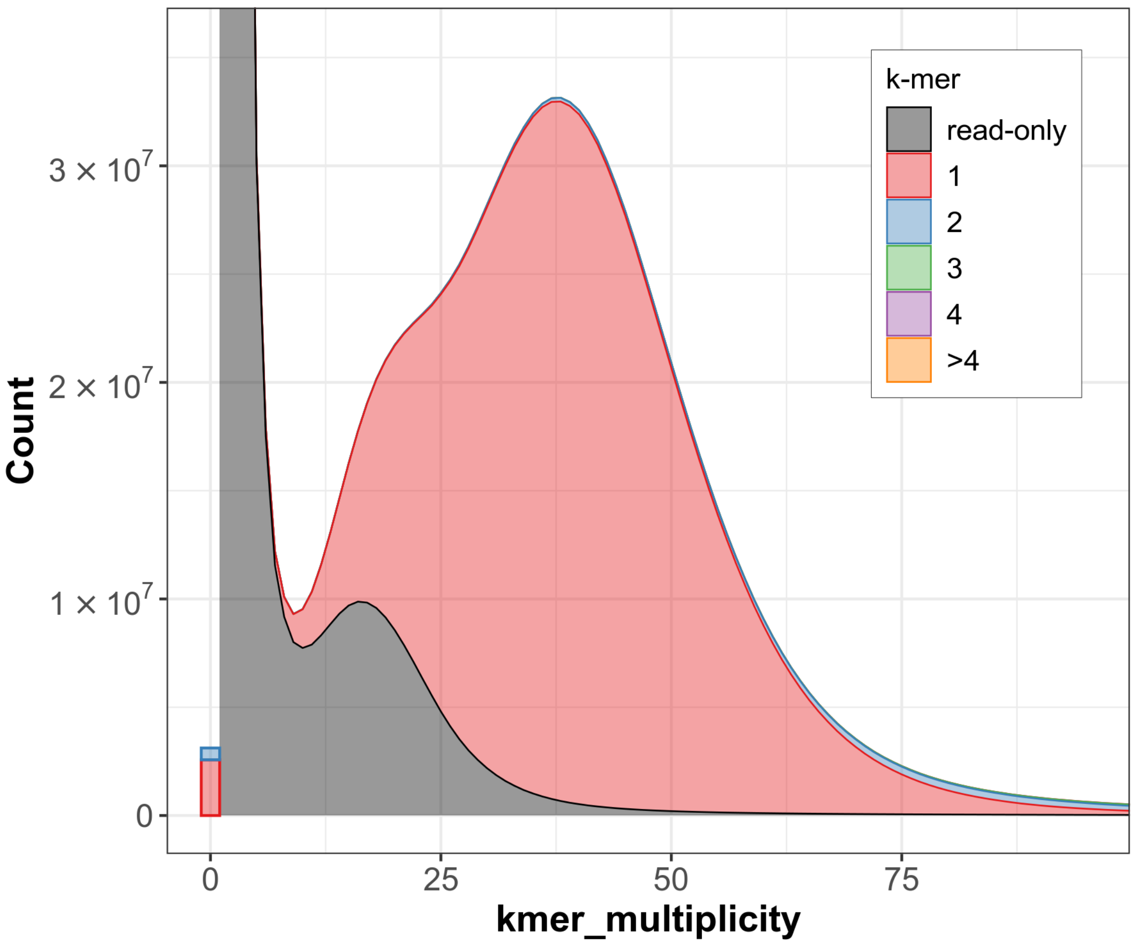


**Figure S3: K-mer comparison between the Illumina reads and the rPodLil1.2 assembly.** Stacked histogram of k-mer distributions obtained by comparing the assembly with Merqury v1.1 using k=21 on the 10X Illumina reads. Artificial duplications corresponding to duplicate k-mers are shown in blue above the main peak (~40x). They only account for 0.68% of the k-mers.

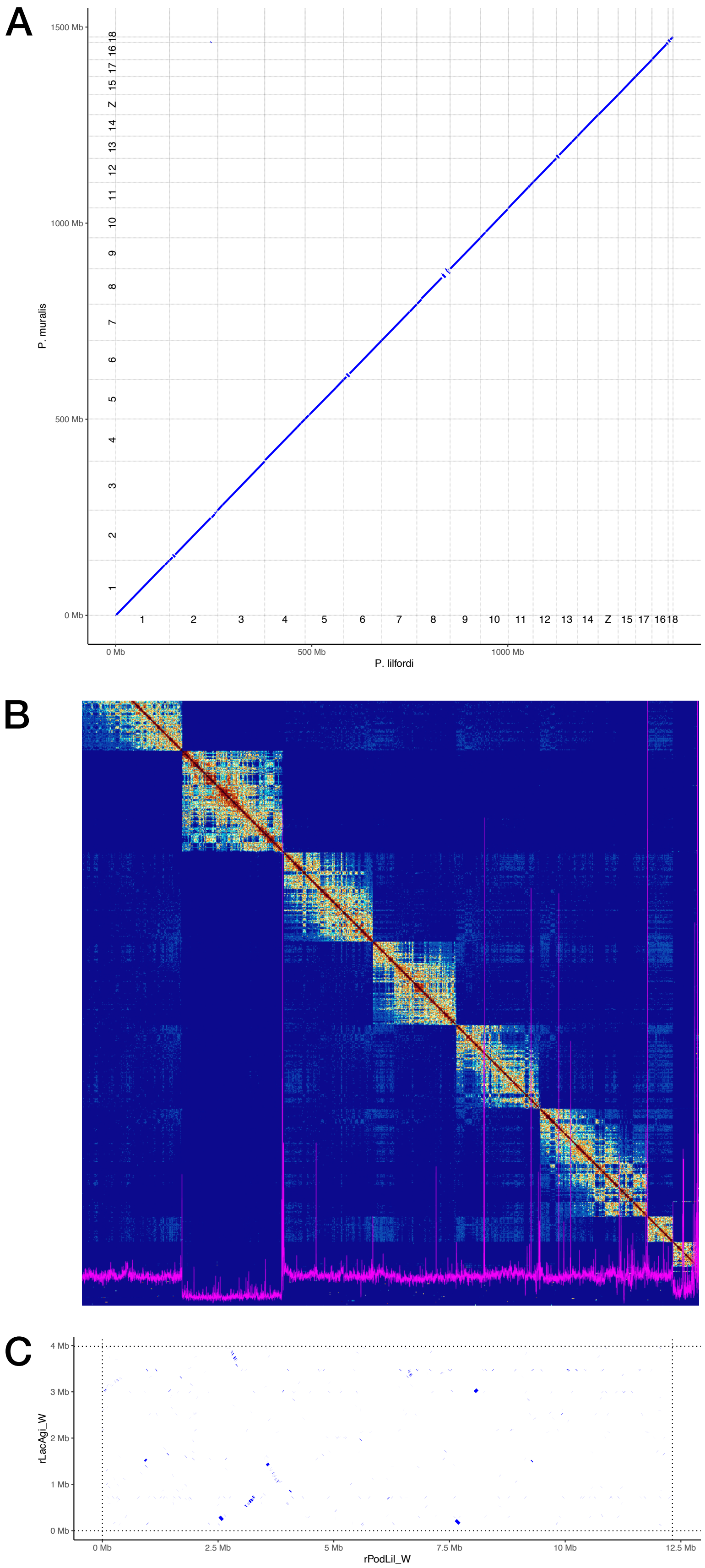


**Figure S4: Chromosome assignment.** (A) Whole genome alignment of *P. lilfordi* to *P. muralis*. Chromosomal sequences, named according to corresponding chromosomal sequences in *P. muralis*, are ordered from largest to smallest in *P. lilfordi* and oriented with respect to *P. muralis*, which inverts the order of chromosomes 16 and 17. Alignments longer than 100 kb were selected and visualised with the *pafr* R library. (B) Hi-C contact map showing scaffolds corresponding to the sexual chromosomes. (C) Alignment of the scaffold assigned to the W chromosome in *L. agilis* against the corresponding scaffold in *P. lilfordi.*
